# Crackling Cloud: an event-driven, cloud-based CRISPR-Cas9 guide RNA design tool

**DOI:** 10.1101/2024.12.04.626718

**Authors:** Jacob Bradford, Divya Joy, Mattias Winsen, Nicholas Meurant, Mackenzie Wilkins, Laurence Wilson, Denis Bauer, Dimitri Perrin

## Abstract

Gene editing has been revolutionised by the CRISPR-Cas9 technology. The versatility and ease-of-use of the technology far exceeds its predecessors, however, the selection of a high-quality guide RNA (gRNA) is critical to directing it to a target site. Selecting gRNA calls upon high-performance algorithms that evaluate nuclease activity at on-target and off-target sites. While there are a suite of programs available, many struggle to analyse the largest genomes, or their predictive accuracy is low. We have previously published a program, named Crackling that is amongst the fastest and most accurate tools available, however, it requires an end-user to have access to a traditional high-performance computing environment. Here, we present an adaptation of Crackling, named Crackling Cloud, that takes advantage of modern serverless cloud technologies that are widely available to anyone, and do not consume resources and incur costs when sitting idle, but can scale to use large volumes of resources when analyses require that. Crackling Cloud is provided as a templated solution using technologies of Amazon Web Services, and is available for free on GitHub under the terms of the BSD 3-clause licence: https://github.com/bmds-lab/Crackling-AWS

## Introduction

CRISPR-Cas9 has become the gold-standard technology for gene editing due to its simplicity and low cost. When a CRISPR-Cas9 nuclease is provided with a guide-RNA (gRNA), it can introduce a double-stranded DNA break at nearly any genomic loci of interest, giving the opportunity for that DNA site to be edited with great precision (1). It has truly modernised the gene editing toolset and has enabled large-scale and robust genomic studies. However, that has only been achieved through the design of high-quality gRNAs using high-performance computing.

There are two considerations when choosing a good-quality gRNA: (1) the on-target activity of the gRNA, often referred to as gRNA efficiency, and (2) the off-target activity of the gRNA, often referred to as gRNA specificity. A good quality gRNA is optimised such that on-target activity is maximised and off-target activity is minimised.

We have optimised the selection of efficient gRNA by using a consensus-based method: we deem gRNA to be efficient when at least two of three independent methods agree (2). That approach is more precise than any individual method alone and has resulted in our experiments seeing successful edits up to 99% of the time (3).

We have also carefully considered how the specificity of gRNA is evaluated. A naive approach would compare each gRNA to every CRISPR targetable site within the genome. For large genomes, like plants and mammals, there can be hundreds of thousands of candidate gRNA, and hundreds of millions of CRISPR sites. Therefore, requiring trillions of sequence comparisons to evaluate gRNA specificity. Instead, for a given gRNA, our method extracts more closely related CRISPR sites by considering its composition of 4-mers (4). Our previously published method for designing good quality gRNA, named Crackling, is amongst the fastest tools available, and uses these improved selection methods (4).

Crackling is fast and accurate, but not always do end-users have the necessary resources to run the program, and they may not necessarily have the expertise to work with bioinformatics programs. In an effort to overcome that, we are turning to high-performance computing environments provided by Amazon Web Services so users can click-to-deploy the software in their own cloud environment. Specifically, we are leveraging the dynamic compute environments available in the suite of “serverless cloud computing” technologies.

Serverless cloud computing refers to a service model that abstracts the underlying compute system away from the developer, and has been demonstrated as a viable platform for omics projects (5). Rather than the research effort involving configuring and spinning up compute clusters to run experiments, efforts are devoted to actual science: designing methods for selecting good quality gRNA. After having those methods in place as software programs, they can be provided to a cloud vendor to execute upon events occurring, such as, an end-user requesting gRNA for a particular gene of a specified species (e.g., HSF1 in acropora millepora, as we have (3)). When no analysis is running, the infrastructure scales down to zero, releasing resources to other customers of the vendor, and incurring no idle cost, unlike traditional server-based architectures.

In this work, we demonstrate an adaptation of Crackling that uses the serverless cloud computing model. That has increased: accessibility to Crackling by removing the need for specialist hardware resources and software skills to run the standalone version; scalability of Crackling by leveraging the seemingly-infinite resources available in the cloud; and, data privacy by enabling end-users to bring the software to their data rather than their data being sent to the software.

To that end, we present Crackling-Cloud: the first free, serverless-cloud-based CRISPR gRNA design pipeline. Crackling Cloud is publicly available on GitHub as opensource software under the terms of the BSD 3-clause licence: https://github.com/bmds-lab/Crackling-AWS

## Methods

Crackling-Cloud is an event-driven pipeline that leverages a flexible, serverless architecture. The architecture could be implemented using the technologies of most modern cloud vendors. We selected Amazon Web Services (AWS) based on their position as the leading vendor and their robust service offering. AWS provides cloud services to many of the largest companies in the world, and to many research and education institutions. To that end, this section describes the implementation of Crackling-Cloud. See Figure 1 for an architecture diagram.

**Fig. 1.**
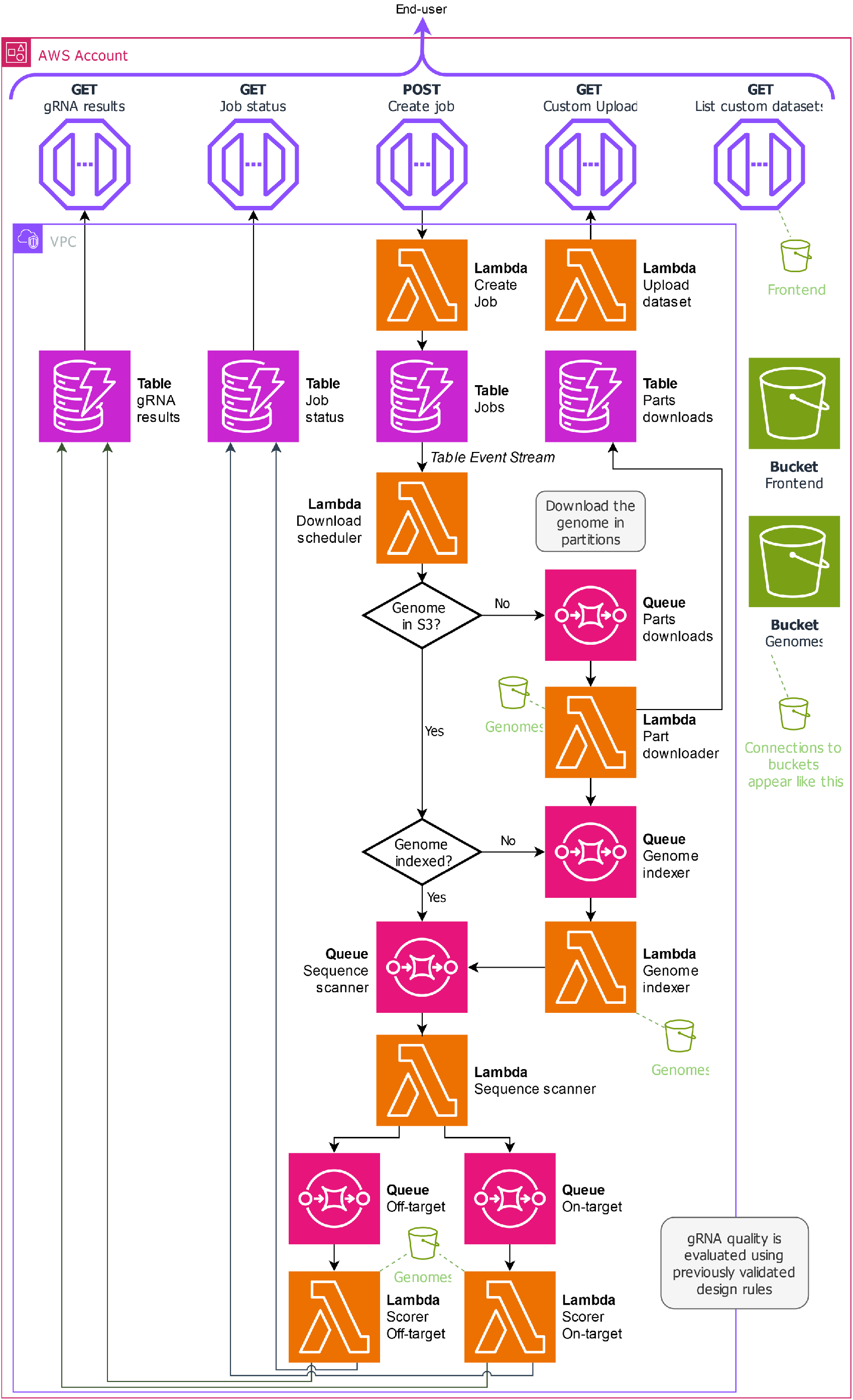
Architecture diagram. Each block represents a cloud service. Lavender hexagons indicate API endpoints, orange lambda symbols indicate function-based compute, purple lightning bolts indicate database tables, green buckets indicate object storage, and magenta circles indicate message queues. Arrows between blocks show data flow. Some services are deployed within a Virtual Private Cloud (VPC) to enhance security, while all services operate within a designated cloud account. The gRNA service is accessible via an HTTP API or the accompanying web client.

The entry point to the pipeline is via a HTTP API served through Amazon API Gateway. There are two critical endpoints that enable an end-user to design gRNA: submit job and retrieve results. To submit a job, the end-user provides the sequence of the gene to target, and to measure gRNA specificity across the entire genome, they provide a NCBI Genome identifier. In return, a job identifier (ID) is provided. Upon submitting a job, the analysis infrastructure spins up from zero. The volume of needed resources is determined by the number of gRNA to assess. Jobs are submitted in series but run in parallel.

If the user wishes to design gRNA for a genome not available in the National Center for Biotechnology Information (NCBI) Genome database, they can securely upload their own into a private Amazon Simple Storage Service (S3) bucket. S3 is an object storage service, akin to a conventional file system.

Results are associated with the job ID and can be retrieved by querying the second API endpoint or through a provided web-based interface. Alternatively, the HTTP API enables an advanced user to retrieve results using their own method (e.g., a custom-built software script, their own graphical interface, Excel, etc.).

Upon a job being submitted via a HTTP request, a database record is created in Amazon DynamoDB. The event of creating that record triggers further data preparation steps, orchestrated by AWS Lambda and Amazon Simple Queue Service (SQS).

Lambda is a serverless run-time environment: no provisioning of servers is required; AWS handles that in the background as needed. Only the code and any dependencies not installed by AWS are provided by the developer. SQS is a queuing service that triggers Lambda to automatically handle, process and scale tasks based on the queued workload.

Lambda can receive messages (tasks) from a SQS queue in batches. Each Lambda invocation processes batches of messages for up to 15 minutes, and without exceeding the 10 gigabyte memory limit. We have configured the size of batching for each SQS-to-Lambda integration based on the specific requirements of the task. When the queue exceeds the current processing capacity, Lambda automatically handles increasing concurrency to handle the workload.

Once a job is submitted to the database, a Lambda function checks that the specified genome is available in storage. If it is not, the NCBI Genome database is queried using the provided accession to retrieve metadata, including the size of the genome. For genomes larger than 50 megabytes, the download is divided into portions, with each portion queued as a byte range of the original file. This approach allows the genome download to be parallelised across multiple Lambda invocations, improving efficiency and avoiding the 15 minute time-limit of Lambda. A subsequent Lambda function handles downloading each portion to S3. After all portions have been obtained, S3 merges the portions.

After merging, an index of all CRISPR-targetable sites is generated. This is critical for evaluating the off-target risk of candidate gRNA. To index the sites, a specialist data structure, named Inverted Signature Slice Lists, is used, as implemented in the standalone edition of Crackling (4). The task of generating the ISSL index is queued and handled by a specialist Lambda function.

Upon the index becoming available, a subsequent Lambda function processes the provided gene sequence. Candidate gRNA are extracted using a regular expression pattern that matches the conventional SpCas9 Protospacer-Adjacent Motif (i.e., 21 nucleotides followed by GG, and the complement for the reverse strand). Each candidate gRNA is added to two separate queues: one for assessing on-target activity and another for assessing off-target activity. The tasks in those queues are consumed by Lambda functions respective to the queue’s purpose.

Due to the asynchronous design, a third API endpoint is available to obtain a progress update. Upon the evaluation of each gRNA property, results are written to the database and can be retrieved via the API or the provided web client.

## Results

We have previously benchmarked the performance of CRISPR-Cas9 gRNA design tools, in terms of speed and accuracy (4, 6). Crackling is amongst the fastest and most accurate tools available. Here, we have benchmarked Crackling Cloud in terms of speed, granted that the gRNA selection process remains the same as the standalone tool, and therefore, its accuracy does not change.

Run-time was measured from when the job was submitted via the API to the completed assessment of every gRNA. That included the time to download and process the genome, which is unlike the previous benchmark, that only included gRNA evaluation and excluded time spent downloading and indexing the genome. Our intent was to measure the actual time that an end-user would wait for all results to become available.

Each experiment was performed as a *warm* run, meaning the infrastructure had already been initialised and was standing by, ready for processing, before timing out and scaling back to zero. In contrast, a *cold* run would have involved additional overhead from launching the serverless architecture, resulting in longer execution times—a typical characteristic and side effect of the serverless cloud computing model. The duration of a cold start can vary from a few milliseconds to over a second.

### Increasing number of gRNA to assess

To measure the impact of the number of gRNA to assess, we generated artificial gene sequences containing a precise number of gRNA. We used these gene sequences and the O. sativa (GCF_001433935.1) genome as inputs. Crackling Cloud analysed 1000 gRNA in 69 seconds, and 10,000 gRNA within 3.5 minutes. See Table 1 for all results.

**Table 1.** Time taken to process the O. sativa genome with an increasing the number of gRNA to assess.

| Number of gRNA | Run-time (seconds) |
| --- | --- |
| 1000 | 69 |
| 2000 | 84 |
| 3000 | 102 |
| 4000 | 117 |
| 5000 | 152 |
| 6000 | 176 |
| 7000 | 170 |
| 8000 | 178 |
| 9000 | 195 |
| 10000 | 209 |

### Increasing genome size

To measure the impact of genome size, we analysed genomes of varying size from the NCBI Genome database. The sequence of the *rrs* gene (16S ribosomal RNA; NCBI gene ID 2700429) was used for each test. rrs was selected as representation of any real sequence that may be provided. It contains 260 candidate gRNA. See Table 2 for results.

**Table 2.** Time taken to process the rrs gene sequence with an increasing genome size.

| Genome | Accession number | Size, Mb | Off-target sites count | Run-time |
| --- | --- | --- | --- | --- |
| *Oryza sativa* | GCF_001433935.1 | 374.4 | $61.7 * 10^6$ | 420 |
| *Panicum hallii* | GCF_002211085.1 | 507.4 | $91.9 * 10^6$ | 667 |
| *Xiphophorus couchianus* | GCF_001444195.1 | 685.5 | $100.6 * 10^6$ | 694 |

## Discussion

### gRNA design is computationally demanding

The design of gRNA is computationally challenging, as the task quickly becomes more expensive proportional to genome size. We previously benchmarked gRNA design tools and found that many cannot scale to large genomes (6). Some tools, in an attempt to overcome that, call upon specialist hardware and compute resources such as on-premises HPC clusters, but those resources may not be readily available to end-users of these tools. Whereas, cloud offers highly scalable and available systems to anyone. Crackling Cloud takes advantage of the seemingly-infinite scale of the cloud.

### Bioinformatic skillsets

gRNA design tools are needed by end-users that may not be equipped with the technical knowledge to install and run bioinformatic tools. Crackling Cloud simplifies the set-up process greatly. Users can quickly and easily deploy the application to their AWS environment using the AWS-native template which describes the needed infrastructure and how to install the software. The end-user requires only an AWS account (simply username, password and a credit card for any costs incurred, which are minimal). Users are not required to configure or manage servers, unlike traditional cloud-based solutions. Once deployed, a webinterface is used to start analyses and obtain results, or advanced users can query the database via the HTTP API.

### Data privacy

While some gRNA design methods are obtained from source and are, therefore, difficult to install, some are offered as “web servers” (in bioinformatics, web servers broadly refer to a website running on a traditional server that provides an interface to a bioinformatic tool installed on the same machine). The gRNA design tools in that category require end users to upload and share data with a third-party, which raises concerns around data sovereignty and organisational constraints. By implementing Crackling Cloud as a cloud-native pipeline that is available as a template, users can deploy a gRNA design tool in their own cloud environment. Now, they can bring the software to their data, rather than sending their data to the software. They can, if so desired, implement their own user access management and security measures, on top of the ones we provide by default. This provides control over where and how their data is stored and shared, ensuring data privacy.

### Serverless model

Traditional server-based architectures are useful for large and long-running tasks. If the average capacity across a cluster is kept to a maximum, the approach becomes economically sound. However, the reality is that in the research context, that is often not the case. Having servers sit idle for long periods is wasteful and costly. The serverless model defers the impact of that by releasing the resources to other customers of the cloud vendor, incurring no idle cost when there is no work to complete. Furthermore, the friction of software development is reduced as the developer no longer needs to provision and maintain server infrastructure. Crackling Cloud uses the serverless model to reduce idle cost to nearly zero by scaling down when there are no gRNA to design.

### Templated cloud architecture

Crackling Cloud is built using the AWS Cloud Development Kit, which provides pre-secured, purpose-built resources designed to be integrated. While that decreased software development efforts, it increased the robustness and security of the software, and makes Crackling Cloud available as a templated solution for any AWS account. We have provided documentation to assist future development efforts, including how to replace gRNA design rules in the relevant analysis components of the architecture.

## Conclusion

Crackling Cloud is a modern CRISPR-Cas9 gRNA design tool that leverages the highly available and scalable technologies available in the cloud. By providing a templated solution, anyone can deploy the software into their own cloud environment. The event-driven architecture keeps development and maintenance efforts low, while also significantly reducing cost when the software is sitting idle. All together, this solves some of the challenges with already available gRNA design tools that are limited to high-performance computing environments and the complementary skills to use. Crackling Cloud is available under the terms of the BSD 3-clause licence and is found on GitHub at https://github.com/bmds-lab/Crackling-AWS

## ACKNOWLEDGEMENTS

We would like to thank Diego Ocando Quintero for his assistance in the development of this research as a student of Queensland University of Technology. This work is supported in part by funds to J.B. from the Commonwealth Scientific and Industrial Research Organisation (CSIRO). D.P. is supported by the Australian Research Council (ARC Discovery Project DP210103401).

## Notes

### Competing Interest Statement

The authors have declared no competing interest.

